# VIRify: an integrated detection, annotation and taxonomic classification pipeline using virus-specific protein profile hidden Markov models

**DOI:** 10.1101/2022.08.22.504484

**Authors:** Guillermo Rangel-Pineros, Alexandre Almeida, Martin Beracochea, Ekaterina Sakharova, Manja Marz, Alejandro Reyes Muñoz, Martin Hölzer, Robert D. Finn

## Abstract

The study of viral communities has revealed the enormous diversity and impact these biological entities have on a range of different ecosystems. These observations have sparked widespread interest in developing computational strategies that support the comprehensive characterization of viral communities based on sequencing data. Here we introduce VIRify, a new computational pipeline designed to provide a user-friendly and accurate functional and taxonomic characterization of viral communities. VIRify identifies viral contigs and prophages from metagenomic assemblies and annotates them using a collection of viral profile hidden Markov models (HMMs). These include our manually-curated profile HMMs, which serve as specific taxonomic markers for a wide range of prokaryotic and eukaryotic viral taxa and are thus used to reliably classify viral contigs. We tested VIRify on assemblies from two microbial mock communities and a large metagenomics study. The results showed that VIRify was able to identify sequences from both prokaryotic and eukaryotic viruses, and provided taxonomic classifications from the genus to the family rank with an accuracy of at least 95.5%. In addition, VIRify allowed the detection and taxonomic classification of a range of prokaryotic and eukaryotic viruses present in 243 marine metagenomic assemblies. Overall, we demonstrate that VIRify is a novel and powerful resource that offers an enhanced capability to detect a broad range of viral contigs and taxonomically classify them.

## Introduction

Viruses are the most abundant biological entities inhabiting our planet, with an estimated 10^30^ virus-like particles (VLP) present in the world’s oceans (Bergh et al. 1989). Even though viruses are often acknowledged as disease-causing agents in animals and plants, many studies to date have demonstrated the pivotal role that viruses play in shaping microbial populations, particularly in the case of phages and their bacterial hosts (Braga et al. 2018; Clokie et al. 2011; Stern and Sorek 2011; Weinbauer and Rassoulzadegan 2004). Phages are generally present among microbial populations interacting with their bacterial hosts via either of two distinct life cycles, lytic or lysogenic. During lytic infections, phages inject their genome into a susceptible host cell and take over its molecular machinery to generate new progeny phages that are released following host cell lysis. By contrast, lysogenic infections are characterised by the establishment of a relatively quiescent intracellular estate in which phages are usually referred to as prophages, and remain in such estate until an external signal induces the onset of the lytic cycle (Campbell 2003; Howard-Varona et al. 2017).

The advent of high-throughput sequencing (HTS) technologies prompted the development of metagenomics, a field that has enabled the exploration of natural microbial communities in a culture-independent fashion. Via metagenomics and the use of methods that concentrate and purify viral particles from environmental samples, it has become possible to catalog the genomic diversity of viruses present in nature and gain insights into the roles they play in different ecosystems (Edwards and Rohwer 2005). Early reports on the application of viral metagenomics (viromics) in a variety of aquatic ecosystems and in the human gut revealed a strikingly large extent of uncharacterized viral genetic diversity, collectively referred to as ‘viral dark matter’ (Reyes et al. 2012; Dutilh et al. 2014; Edwards and Rohwer 2005; Hurwitz and One 2013; Reyes et al. 2010; S. Roux et al. 2012). As HTS technologies became more accessible and novel methods for *de novo* genome assembly from metagenomic datasets were devised, an increasing number of studies published during the last decade led to the discovery of vast numbers of uncultivated virus genomes (UViGs) (Gregory et al. 2019; D. Paez-Espino, Eloe-Fadrosh, and Nature 2016; Shkoporov et al. 2019; Simon Roux et al. 2019). Moreover, these studies have been of paramount importance for understanding the role that viruses play in different ecosystems, which is particularly true for oceanic surveys that evidenced the critical role that viruses play in biogeochemical cycling and metabolic modulation of ocean microbes (Breitbart et al. 2018; Simon Roux et al. 2016; Gregory et al. 2019).

A recent review paper of the TARA Oceans project highlighted the urgent need for automated approaches to systematically organise the enormous virus sequencing data beyond the species level (Sunagawa et al. 2020). Several computational tools and resources that offer a solution to this challenge have been developed and made publicly available in the last two decades (Rohwer and Edwards 2002; Meier-Kolthoff and Göker 2017; Aiewsakun et al. 2018). One of the most prominent and currently available tools is VConTACT2, which employs data on proteins shared between phage genomes to build a similarity network that is further analysed to identify clusters of evolutionarily related phages (Jang et al. 2019). VConTACT2 is part of the current suite of analysis tools offered in iVirus, a community resource available via the CyVerse cyberinfrastructure that provides access to viromics tools and public datasets (Bolduc et al. 2017). An alternative approach to the taxonomic classification of viruses involves the use of protein profile HMMs representing clusters of homologous proteins or protein domains, which in turn serve as markers for different viral taxa. Such an approach is followed by ClassiPhage, a tool that employs a collection of profile HMMs that serve as taxonomic markers of the phage families *Myoviridae, Siphoviridae, Podoviridae* and *Inoviridae* (Chibani et al. 2019). Phage classification with ClassiPhage demonstrated agreement with the reference ICTV taxonomy, although the generation and testing of the taxon-specific profile HMMs was solely focused on phages targeting members of the bacterial family *Vibrionaceae* (Chibani et al. 2019). More recently, a set of 31,150 profile HMMs representing proteins predicted in viral genomes from NCBI were used as the basis for a set of Random Forest classifiers aimed at assigning viral taxonomy at the order, family and genus ranks (Moreno-Gallego and Reyes 2021). These classifiers demonstrated very high classification accuracy (≥ 89%) at all the aforementioned ranks, suggesting that these profile HMMs (or ViPhOGs as they were named) could be the foundation for a new method that taxonomically classifies viral genomic sequences present in metagenomic datasets.

Currently, a few pipelines that are still supported allow users to functionally and taxonomically characterise viral contigs assembled from metagenomic datasets, including VirMiner, iVirus and its simplified version available at the Department of Energy’s Systems Biology KnowledgeBase (KBase) (Zheng et al. 2019; Bolduc et al. 2017, 2021). Here we present VIRify, a new computational pipeline that combines the prediction of virus-like sequences based on canonical viral signals and sequence patterns, with the functional and taxonomic annotation of viral contigs based on a comprehensive collection of publicly available profile HMMs, including a set of manually curated ViPhOGs. VIRify is currently available as an independent pipeline to analyse any user-supplied metagenomic assembly. VIRify reduces the number of false-positive predictions using virus-specific protein models, and has been implemented with portability at the forefront of the design. To run VIRify (https://github.com/EBI-Metagenomics/emg-viral-pipeline), the user can choose between its implementation as a Nextflow workflow (Di Tommaso et al. 2017) or using the Common Workflow Language (CWL) and WorkflowHub (Amstutz, Crusoe, Tijanić, Chapman, Chilton, Heuer, Kartashov, Leehr, Ménager, Nedeljkovich, and Others 2016; Goble et al. 2021). Both Workflow Management Systems rely on the same code base and Docker containers and while the Nextflow version is intended for easy installation and execution as an independent pipeline, the CWL version can be directly integrated into already existing platforms such as MGnify (Mitchell et al. 2020). We showcase the application of VIRify based on two viral mock communities mostly comprised of phages (Kleiner, Hooper, and Duerkop 2015) but also eukaryotic viruses (Conceição-Neto et al. 2015), as well as 243 metagenome assemblies of the TARA Oceans project (Gregory et al. 2019).

## Methods

### Manual curation of virus-specific protein HMMs and definition of model-specific bit score thresholds

The ViPhOG database consists of a collection of 31,150 protein profile HMMs that were built using sequences retrieved from all viral genomes available in April 2015 in NCBI’s RefSeq (Moreno-Gallego and Reyes 2021). We manually curated the taxon-specific profile HMMs by searching the complete ViPhOG database against all entries in UniProtKB (February 2019 version). This was achieved using hmmsearch (v3.2.1) and setting a per-sequence reporting evalue threshold of 1.0×10^-3^. The resulting output was analysed using in-house python scripts (https://github.com/EBI-Metagenomics/emg-viral-pipeline/tree/master/bin/models_vs_uniprot_check) to identify which ViPhOGs could be used as taxon-specific markers, hereafter referred to as informative ViPhOGs. These were identified after applying the following steps to the hmmsearch output of each ViPhOG:

1. Reported taxonomy identifiers (taxid) were recorded, along with the highest bit score obtained for each one of them and their associated taxa at the genus, subfamily, family and order ranks.
2. For each one of the target taxa recorded in the previous step, the corresponding highest bit scores were sorted and a bit score range was defined using the lowest and highest of these values.
3. Starting with the target taxa at the genus rank, it was determined whether the query ViPhOG was associated with a single taxon or the bit score range for the best taxon did not overlap with the range obtained for the remaining taxa. If either of these conditions was fulfilled, then the query ViPhOG was selected as informative at the genus rank. Otherwise, the procedure was repeated at the subfamily, family and order ranks, until the query ViPhOG could be selected as informative for any of them. If the query ViPhOG was not assigned to any taxon at the searched ranks, it was considered non-informative.

To leverage the data obtained from the analysis described above and set inclusion bit score thresholds suitable for each model, we defined two bit score thresholds (S1 and S2) for each informative ViPhOG. S1 was defined as the minimum bit score associated with the ViPhOG’s target taxon, whereas S2 was set as the maximum bit score obtained for other viral taxa identified through the previously described procedure. For the cases in which no further taxa were reported at the same rank as the target taxon, only S1 was set for the corresponding informative ViPhOG. Next, we define GA (gathering), TC (trusted cutoff), and NC (noise cutoff) parameters of each informative ViPhOG model. GA thresholds define reliable curated thresholds for family memberships, NC thresholds describe the highest-scoring known false positive, and TC thresholds refer to the score of the lowest-scoring known true positive that is above all known false positives, which reflects a similar strategy to Pfam (Mistry et al. 2021). In addition, each parameter (GA, TC, NC) defines two thresholds for reporting and inclusion scores: the per-sequence threshold (GA_seq_, TC_seq_, NC_seq_) and the per-domain threshold (GA_dom_, TC_dom_, NC_dom_). We used our previously defined bit score thresholds S1 and S2 to set these six parameters specifically for each informative ViPhOG model as follows:

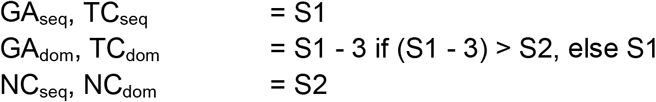

S1 was added as TC and GA per-sequence threshold, and trimmed by three bit to apply a per-domain threshold. Trimming was only applied if S1 did not drop below S2. The S2 bit score was unmodified and used as both per-sequence and per-domain NC. For the cases in which no S2 bit score was recorded, the noise cutoff was omitted and the sequence-specific S1 value was trimmed by three bit to allow less restrictive predictions. Finally, the values set for the GA, TC and NC parameters were added to the header section of each informative ViPhOG’s HMM file.

Viral taxa linked to the informative ViPhOGs were categorized as prokaryotic or eukaryotic based on the host they target. Currently known viral-host relationships were retrieved from Virus-Host DB that comprised 9,616 viral entries targeting eukaryotes and 3,763 viral entries targeting either bacteria or archaea, when it was accessed in November 2019 (Mihara et al., 2016). In addition, we explored the extent to which the informative ViPhOGs covered the different lineages that comprise the current viral taxonomy. Using the ete3 python package (Huerta-Cepas et al., 2016), a cladogram was built including all the viral genera and corresponding ancestral taxa listed in the NCBI Taxonomy database, accessed in March 2020 (Federhen, 2012). The number of informative ViPhOGs identified for each viral genus was mapped on the cladogram using the interactive Tree of Life (iTOL) resource and its associated annotation files (Letunic & Bork, 2019).

### Preparation of mock metagenome assemblies for benchmarking

Paired-end reads of six virus-enriched samples from Kleiner, Hooper, and Duerkop (2015) were downloaded from ENA (study: PRJEB6941; runs: ERR575691, ERR575692, ERR576942, ERR576943, ERR576944, ERR576945) and concatenated to make a single input file. Prior to assembly, host contamination was removed by k-mer-based decontamination against the mouse genome (Ensembl GRCm38, primary assembly) with BBDuK v38.79 from the BBTools suite (https://sourceforge.net/projects/bbmap/) using a k-mer size of 27. To include a metagenome assembly that also comprised eukaryotic viruses, we further obtained paired-end reads of seven virus-enriched samples (study: PRJNA319556; runs: SRR3458563-3458569) from Conceição-Neto et al. (2015).

For the combined data sets within both studies, two individual metagenome assemblies were performed by first cleaning the reads with fastp v0.20.0 (Chen et al. 2018) and passing them to metaSPAdes v3.14 (Nurk et al. 2017) using default parameters. The ‘--only-assembler’ parameter was used to generate the assembly of the combined Neto read sets due to memory constraints. We used QUAST v5.0.2 (Gurevich et al. 2013) to assess basic quality metrics of the assemblies. The two assemblies (hereafter named Kleiner and Neto assemblies) can be downloaded from the Open Science Framework (https://doi.org/10.17605/OSF.IO/FBRXY).

### Selection and comparison of virus prediction tools

We used the multi-tool workflow “What the Phage” (WtP) (Marquet et al. 2020) for comparing different virus prediction tools in an attempt to identify the best tool combination for a comprehensive initial virus prediction of our pipeline. We run WtP release v0.9.0 including VirSorter (with and without virome option), VirFinder (default and VF.modEPV_k8.rda models), PPR-Meta, DeepVirFinder (Ren et al. 2020), MARVEL (Amgarten et al. 2018), metaPhinder (Jurtz et al. 2016), VIBRANT (Kieft, Zhou, and Anantharaman 2020), VIRNET (Abdelkareem et al. 2020), Phigaro (Starikova et al. 2020) and sourmash (Pierce et al. 2019) on the Kleiner and Neto assemblies. The analysis was performed on contigs that were at least 1.5 kb long to filter out shorter contigs with potentially few ORFs, which are less likely to be correctly classified by our ViPhOG-based taxonomic annotation pipeline. Previous studies have revealed that viral genomes shorter than 1.5 kb tend to code for no more than 5 proteins (Moreno-Gallego and Reyes 2021). Based on a previous study in which global oceanic viral populations were surveyed, we manually updated the parameter configuration file of WtP to include all VirSorter predictions from categories 1-5 and to filter VirFinder results by p-values < 0.05 and scores ≥ 0.7 (Gregory et al. 2019).

To assess the performance of each viral prediction tool and the tool combination implemented in VIRify, we identified the viral contigs in the Kleiner and Neto assemblies by aligning them to the genomes of the viruses that comprised the corresponding mock communities. For both mock communities the set of viral genomes used as targets for the alignments included all the viruses listed in the corresponding studies, and the putative prophages predicted in the genomes of the bacteria that were either part of the mock communities or used for propagating the mock community phages. Thus, prophage prediction was conducted for the following bacterial genome sequences: *Pseudomonas savastanoi* pv. *phaseolicola* strain HB10Y (GCF_001294035), *Salmonella enterica* subsp. *enterica* serovar Typhimurium str. LT2 (NC_003197), *Listeria monocytogenes* EGD-e (NC_003210), *Bacteroides thetaiotaomicron* VPI-5482 (NC_004663), *Enterococcus faecalis* V583 (NC_004668), *Escherichia coli* B strain C3029 (NZ_CP014269), *Lactobacillus acidophilus* strain ATCC 4356 (GCF_000786395), *Bifidobacterium animalis* subsp. *lactis* ATCC 27674 (GCF_001263985), *B. thetaiotaomicron* strain ATCC 29741 (GCF_004349615) and *E. coli* DSM 30083 (NZ_CP033092). Prophages were predicted from the listed bacterial genomes using VirSorter (Simon Roux, Enault, et al. 2015), selecting the Viromes database and activating the virome decontamination mode, and PHASTER (Arndt et al. 2016).

VirSorter predictions in categories 4 and 5, and PHASTER predictions classified as Intact or Questionable were clustered using cd-hit-est with a sequence identity cut-off of 0.9 and default settings (Li and Godzik 2006). The set of putative prophages selected for each mock community included all VirSorter category 4 predictions, PHASTER Intact prophages, and regions that were reported both as VirSorter category 5 predictions and PHASTER Questionable prophages. Contigs from each mock community assembly were aligned to the corresponding set of viral genomes using nucmer v3.23 with default settings (Kurtz et al. 2004), and contigs were identified as viral when at least 70% of their length was covered by an alignment with at least 90% sequence identity. Following the identification of viral contigs within the Kleiner and Neto assemblies, the performance of the viral predictions tools tested using WtP and the combination selected for VIRify was measured via the calculation of the F1-score.

The output from WtP was visualised in UpSet plots and presence/absence maps that indicated whether an input contig was predicted as viral or not by each of the assessed tools. MARVEL and VIRNET predictions were not included for the Neto assembly because both tools failed to process this assembly for unidentified technical reasons.

### VIRify taxonomic annotation pipeline description

#### Identification of putative viral sequences

Based on the results obtained with WtP and the procedure previously employed in a global oceanic survey of viral populations, it was determined that VIRify’s detection of putative viral contigs would be carried out with VirFinder, VirSorter and PPR-Meta, while putative prophage detection would be conducted with VirSorter only (**Figure 1**) (Gregory et al. 2019; Ren et al. 2017; Simon Roux, Enault, et al. 2015; Fang et al. 2019). Viral prediction with VirSorter is carried out using parameters ‘--db 2’ and ‘--virome’ when the processed metagenomic assemblies are generated from virome datasets (as was the case for VIRify’s benchmarking with the Kleiner and Neto assemblies), otherwise the latter parameter is not set. Putative prophages are retrieved from VirSorter predictions in prophage categories 4 and 5 as defined by the tool. Detection of putative viral contigs with VirFinder is conducted using the classifiers in the model file VF.modEPV_k8.rda, which was trained using sequences from prokaryotic and eukaryotic viruses (available at https://github.com/jessieren/VirFinder). Predicted viral contigs are classified into two different categories within the VIRify pipeline as follows: contigs reported by VirSorter in categories 1 and 2 are placed in the high-confidence category, whilst the low-confidence category includes contigs that satisfy any of the following conditions:

- Reported by VirFinder with p-value < 0.05 and score ≥ 0.9.
- Reported by VirFinder with p-value < 0.05 and score ≥ 0.7, and also reported by VirSorter in category 3.
- Reported by VirFinder with p-value < 0.05 and score ≥ 0.7, and also reported as phage by PPR-Meta.

**Figure 1:**
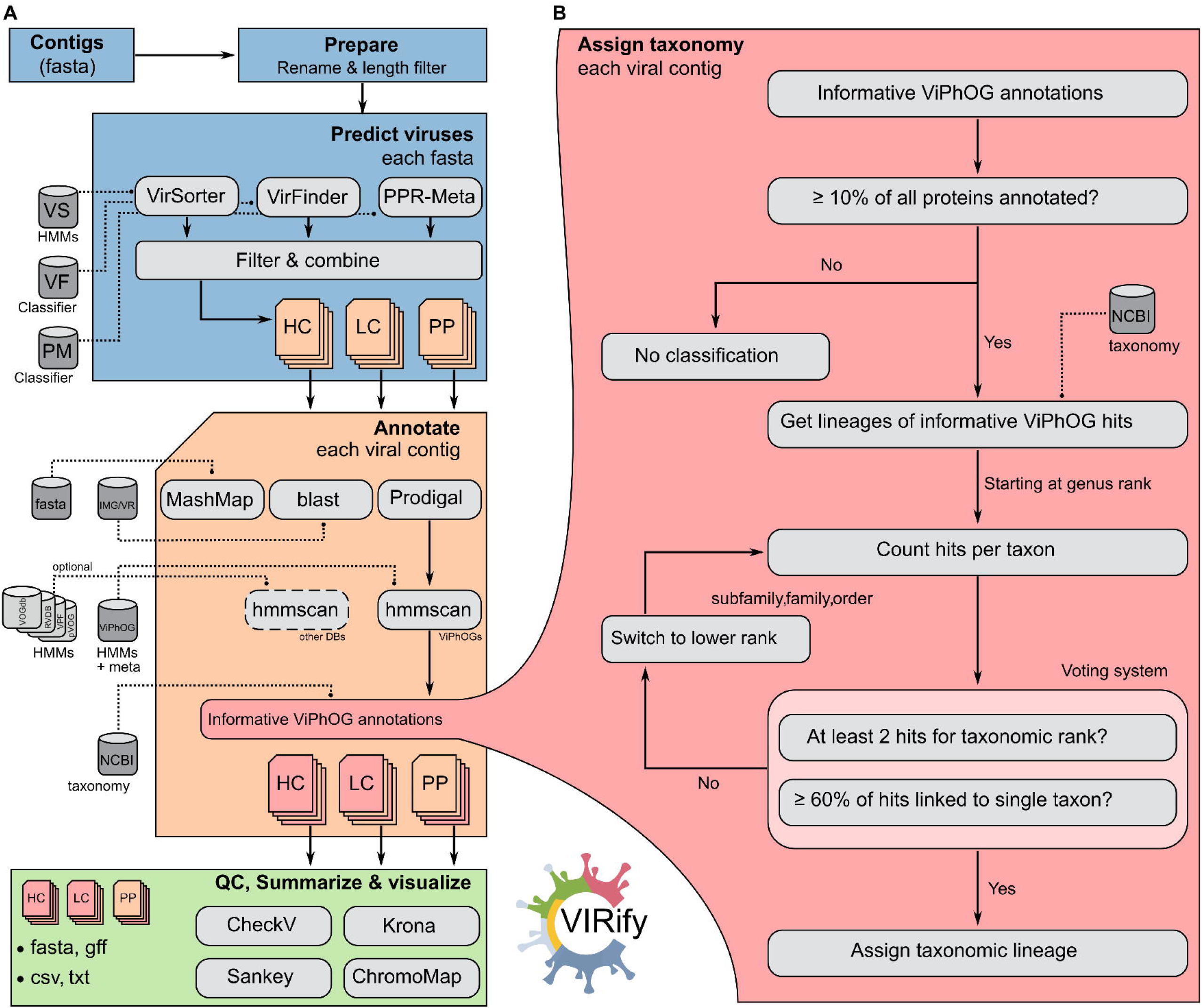
Overview of the VIRify pipeline. **(A)** Starting from a set of contigs (fasta file) the pipeline preprocesses the input sequences (ID renaming, length filtering) and predicts contigs from a putative viral origin that are split into high confidence (HC), low confidence (LC) and putative prophage (PP) sets. Each selected contig is then annotated and assigned to a taxonomy, if possible. All results (annotated viral contigs) are quality-controlled with CheckV, and finally summarised and visualised. **(B)** The assigned taxonomy is based on the informative ViPhOG hits per contig and performed on genus, family, subfamily and finally order rank. We consider high confidence and low confidence ViPhOG hits and discard non-informative models where no clear taxonomy signal could be assigned.

Finally, CheckV (Nayfach et al. 2021) is employed to assess the quality and completeness of the categorised viral genome contigs.

#### Annotation of putative viral sequences with informative ViPhOGs

Protein-coding sequences within putative viral contigs and prophages are detected with Prodigal v2.6.3 (Hyatt et al. 2010), using the standard bacterial/archaeal translation table (-g 11) and the metagenomic prediction mode (-p meta) (**Figure 1**). A recent study in which the performance of a range of ORF-prediction tools was assessed for the viral genomes available in NCBI’s RefSeq reported that Prodigal was the most accurate tool for DNA viruses, which account for ~97% of the viral genomic sequences currently available in public databases (González-Tortuero et al. 2021; Simon Roux et al. 2021) Amino acid sequences derived from the predicted coding sequences are scanned with the complete ViPhOG database using hmmscan v3.2.1 (Finn, Clements, and Eddy 2011) and the model-specific bit score thresholds defined as previously described, in order to supersede any thresholding based on statistical significance alone. To apply the defined model-specific thresholds we set the --cut_ga option during execution of the hmmscan command. However, to account for the set of ViPhOGs for which no bit score thresholds were defined (and therefore set to 0), an additional post-processing filter is applied to the full-sequence e-values reported in the hmmscan output (E ≤ 1.0×10^-3^).

#### Taxonomic assignment of putative viral sequences

In-house python scripts (see https://github.com/EBI-Metagenomics/emg-viral-pipeline) are employed for parsing the hmmscan results and to provide a taxonomic assignment for all putative viral contigs and prophages, based on the reported hits against the ViPhOG database. To guarantee that taxonomic assignments are based on an appropriate amount of evidence, only contigs with at least 10% of encoded proteins having informative hits against the ViPhOG database are considered. Contigs that pass this filtering step are subjected to a voting system that assigns a likely taxonomy to them, based on all corresponding reported hits against the set of informative ViPhOGs. Starting at the genus level, the voting system checks all taxa associated with the reported informative ViPhOGs and provides a genus-level assignment if at least 2 hits are reported at this level and 60% or more of these correspond to the same viral genus. If the first condition is not satisfied the voting system does not assign a taxon at the genus level and continues the analysis at the next taxonomic level up, i.e. subfamily. Alternatively, if the first condition is satisfied but none of the proportions calculated for the reported genera passes the 60% threshold, then the largest proportion value is provided to the user. This process is subsequently repeated at the subfamily, family, and order levels for all contigs that pass the initial filtering step (**Figure 1**). Finally, we assign a virus taxonomy to the putative viral contigs and prophages if the voting criteria are fulfilled. The aforementioned thresholds for the number of informative ViPhOGs used for taxonomic annotation were selected because those values led to the largest number of correct taxonomic assignments for the mock community assemblies.

#### Visualization of assigned viral taxonomies and ViPhOG hits

Putative viral contigs with and without an assigned taxonomy are visualized using interactive Krona (Ondov, Bergman, and Phillippy 2011) and Sankey plots inspired by the Pavian package (Breitwieser and Salzberg 2016). ORFs identified with Prodigal and their corresponding ViPhOG hits are visualized for each contig using the ChromoMap package v0.2 (Anand and Rodriguez Lopez 2020). The package does not visualize exact start and stop positions for each ORF but instead relies on a more general grid view thus visualizing the general coverage of annotated ORFs for each contig.

### Analysis of 243 TARA Oceans assemblies with VIRify

We obtained 243 ocean microbiome assemblies from ENA (https://www.ebi.ac.uk/ena/browser/view/PRJEB22493), generated as part of the TARA Oceans microbiome study (Gregory et al. 2019). The FASTA identifiers include the size fraction of the corresponding sample, thus samples enriched for viruses are labelled with the suffix _0.1-0.22. All assemblies were filtered to retain contigs that were at least 5 kb long and these were screened for viruses using VIRify and setting the --virome option to activate VirSorter’s virome decontamination mode.

### Selection and comparison of virus-specific protein profile HMM databases

The taxonomic assignment of VIRify relies purely on our own protein profile HMM database, consisting of 31,150 so-called ViPhOGs (Leonardo Moreno-Gallego and Reyes 2021), their taxonomy, and model-specific bit score cutoffs. In an effort to be comprehensive, our pipeline also provides the option of additional annotations by incorporating 25,399 models from the VOGdb (http://vogdb.org/), 25,281 viral protein family (VPF) models from IMG/VR (David Paez-Espino et al. 2017, 2019), 9,911 models from RVDB v17.0 (Goodacre et al. 2018; Bigot et al. 2019), and 9,518 models from pVOG (Grazziotin, Koonin, and Kristensen 2017). VIRify can use all databases to individually annotate proteins from all putative viral contigs, using hmmscan with default values and an e-value cutoff of 1.0×10^-3^. However, the final taxonomic assignment provided for the putative viral contigs relies exclusively on our curated set of ViPhOGs.

To compare the annotations provided by all five databases, we ran hmmscan on all predicted proteins from putative viral contigs detected in the two mock community assemblies (Neto and Kleiner) and the 243 TARA Oceans assemblies. To allow for a better comparison against the other databases, we distinguished hits against the ViPhOG models into hits only based on an evalue cutoff of 1.0×10^-3^ and hits additionally identified using our predefined model-specific bit score thresholds.

### General implementation of VIRify

VIRify is implemented using the workflow management systems CWL (Amstutz, Crusoe, Tijanić, Chapman, Chilton, Heuer, Kartashov, Leehr, Ménager, Nedeljkovich, Scales, et al. 2016) and Nextflow (Di Tommaso et al. 2017), and an overview of the pipeline is given in **Figure 1**. All third-party tools are encapsulated in software containers to allow easy distribution of the pipeline on local, cluster, or cloud systems using either CWL or Nextflow. Custom Python scripts connect the output and input of the tools used and are available via GitHub (https://github.com/EBI-Metagenomics/emg-viral-pipeline/tree/master/bin). Both workflow implementations rely on the same scripts and software containers. With configured Nextflow and Docker (Merkel and Others 2014) installations, VIRify can be simply downloaded and run with a single command:

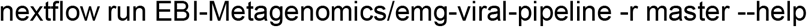

All required databases and metadata files are automatically downloaded and stored for later (offline) reuse. We recommend running a stable release version from the repository which can be selected via the -r flag. Different Nextflow profiles allow the execution on a local system or cluster (currently supported are LSF and SLURM). The pipeline may also be run in an “annotation” mode that skips the prediction of putative virus sequences and directly assigns viral taxonomies to all the input contigs.

## Results

### Viral diversity is comprehensively covered by the set of manually curated informative ViPhOGs

The ViPhOG database consists of 31,150 profile HMMs that were created using proteins from viral genomes found in NCBI’s RefSeq database (Moreno-Gallego and Reyes 2021). Based on the search of homologous sequences in the UniProtKB database and the subsequent analysis conducted on the output data, we identified 22,013 informative ViPhOGs that could serve as viral taxon-specific markers. **Figure 2** indicates the number of informative ViPhOGs obtained for each of the analysed taxonomic ranks (genus, subfamily, family and order), and for each rank it illustrates the proportion of targeted taxa that correspond to either prokaryotic or eukaryotic viruses. Overall, 78.6% of the informative ViPhOGs were linked to taxa at the genus level and the numbers identified for the remaining ranks decreased from the subfamily to the order level. Furthermore, a virtually equal proportion of informative ViPhOGs was identified for prokaryotic and eukaryotic viruses (50.7% for the former and 49.3% for the latter). Regarding the coverage of currently known viral taxa, the informative ViPhOGs cover 48.1, 72.5, 33.8 and 57.1% of genera, subfamilies, families and orders represented in the viral NCBI taxonomy retrieved in March 2020 (**SFigure 1**). As illustrated in the figure, our set of informative ViPhOGs includes taxonomy markers for many representative viral lineages such as the ubiquitous and widely-described order *Caudovirales*, the eukaryotic virus order *Herpesvirales*, and the eukaryotic viral families *Mimiviridae*, *Coronaviridae*, and *Poxviridae*, among others. Moreover, based on the coverage of the NCBI viral taxonomy of March 2020, only 10.7% of viral genera had no informative ViPhOGs associated with any of the taxa that comprise their corresponding lineages. Thus, our set of informative ViPhOGs cover 89.3% of all the lineages represented by the currently known viral diversity in at least one taxonomic level (**SFigure 1**).

**Figure 2:**
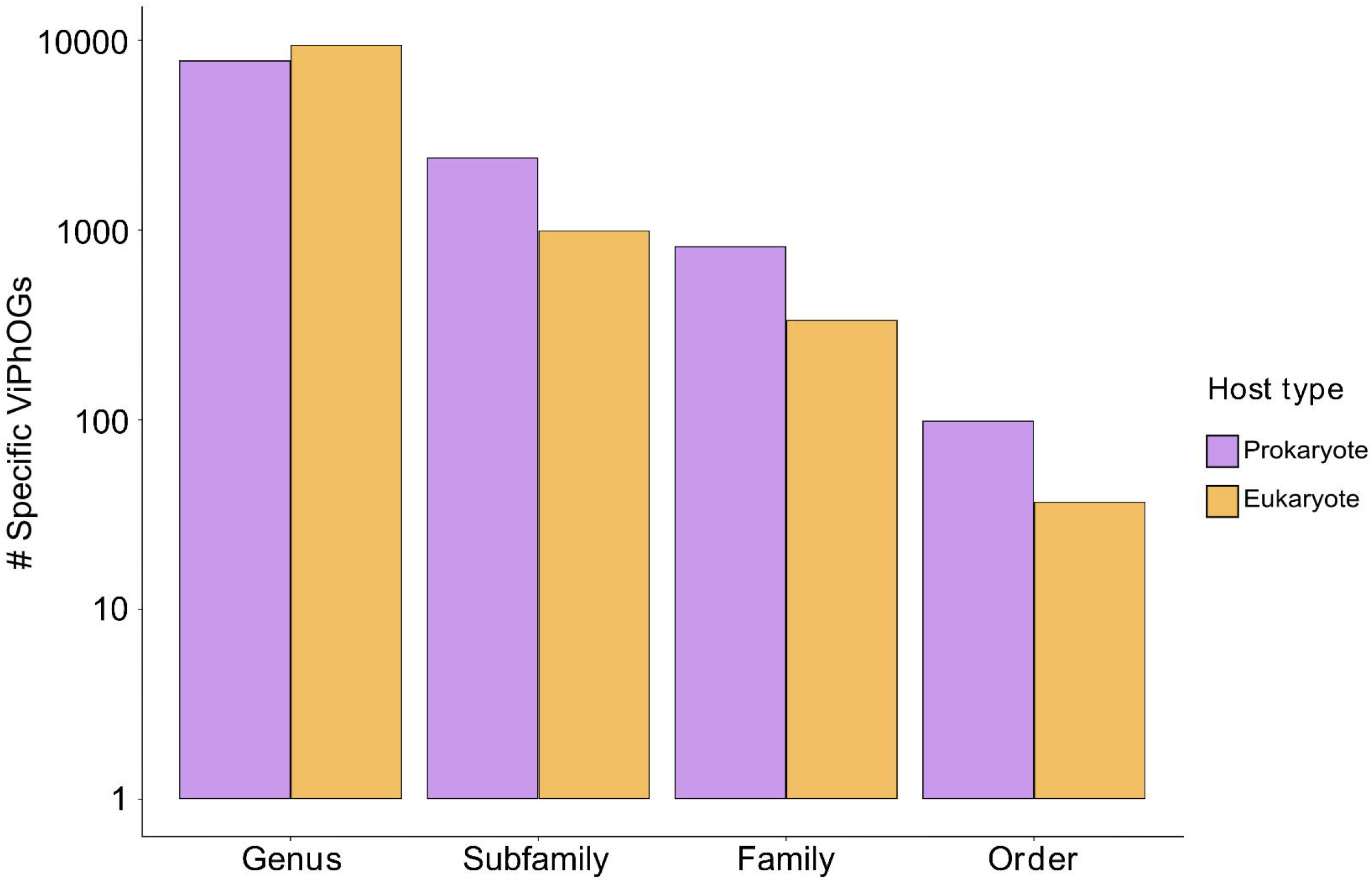
Number of specific ViPhOGs identified for different viral taxonomic ranks. 31,150 ViPhOGs were searched against all entries in UniProtKB, and based on the output they were designated as specific for different viral taxa (see Methods). Purple refers to specific ViPhOGs assigned to prokaryotic viral taxa, whereas yellow indicates specific ViPhOGs for eukaryotic viral taxa.

### VIRify accurately classifies viruses on co-assemblies of two largely viral mock communities

VIRify was tested on datasets generated from viral mock communities to evaluate its viral sequence prediction performance and compare its taxonomic assignments with the taxonomy of the viruses present in the communities. The selected datasets were generated for two different studies that evaluated a range of viral enrichment and purification procedures that were applied to mock microbial communities. Kleiner et al. (2015) sequenced a mock microbiome prepared in mouse faeces using bacterial and phage cultures (germ-free C57BL/6 J mice). The samples contained six different phages in varying concentrations: P22, T3, T7, □6, M13 and □VPE25. Neto et al. (2015) constructed a mock microbiome comprised of four common gut bacteria and nine highly diverse viruses, most of which represent the eukaryotic viral families *Circoviridae, Parvoviridae, Polyomaviridae, Alphaflexiviridae, Reoviridae, Coronaviridae, Herpesviridae* and *Mimiviridae*, and including a phage from the family *Ackermannviridae* (for details see Table 1 in Conceição-Neto et al. 2015). The nine viruses differ highly in genome length (1.8□kb to 1,180□kb), genome type (dsDNA, dsRNA, ssDNA, ssRNA), and genome compositions (linear, circular, segmented). Co-assemblies of datasets obtained from viral-enriched samples were generated for each of the selected studies and used as input for the VIRify pipeline. The Kleiner co-assembly comprises 5,310 contigs of which 224 contigs have a length >= 1.5 kb with an N50 of 74,309 bp. The Neto co-assembly is composed of 341,587 contigs of which 1,686 are >= 1.5 kb with an N50 of 16,751 bp. Thus, 95.78% of the contigs in the Kleiner assembly, and 99.51% of the contigs in the Neto assembly are shorter than 1500 bp.

**Table 1.** VIRify’s viral prediction results for two mock community assemblies. The analyses were conducted for all assembled contigs longer than 1.5 kb.

| Assembly | Total contigs | Viral contigs | True positives | True negatives | False positives | False negatives |
| --- | --- | --- | --- | --- | --- | --- |
| Kleiner | 224 | 17 | 13 | 206 | 1 | 4 |
| Neto | 1686 | 259 | 134 | 1390 | 37 | 125 |

#### VIRify comprehensively selects putative viral contigs

We ran a modified version of WtP (see Methods) on both mock community assemblies (Kleiner, Neto) to compare ten different tools for virus prediction. **SFigure 2** illustrates the performance of all the assessed tools for each of the analysed mock community assemblies. Our results show that a selection of VirSorter, VirFinder, and PPR-Meta sufficiently represented a common proportion of putative viral contigs from both mock community assemblies across all compared prediction tools (**Figure 3A**, **SFigure 2** and **SFigure 3**). Thus, the combination of these tools was selected as VIRify’s method for viral contig prediction, although the selection of putative viral contigs from this combination is subjected to a set of rules similar to the selection criteria used in a previous study of global oceanic viral populations (see Methods section for further details) (Gregory et al. 2019). **Table 1** summarises the viral contig prediction results obtained with VIRify for the Kleiner and Neto assemblies.

**Figure 3:**
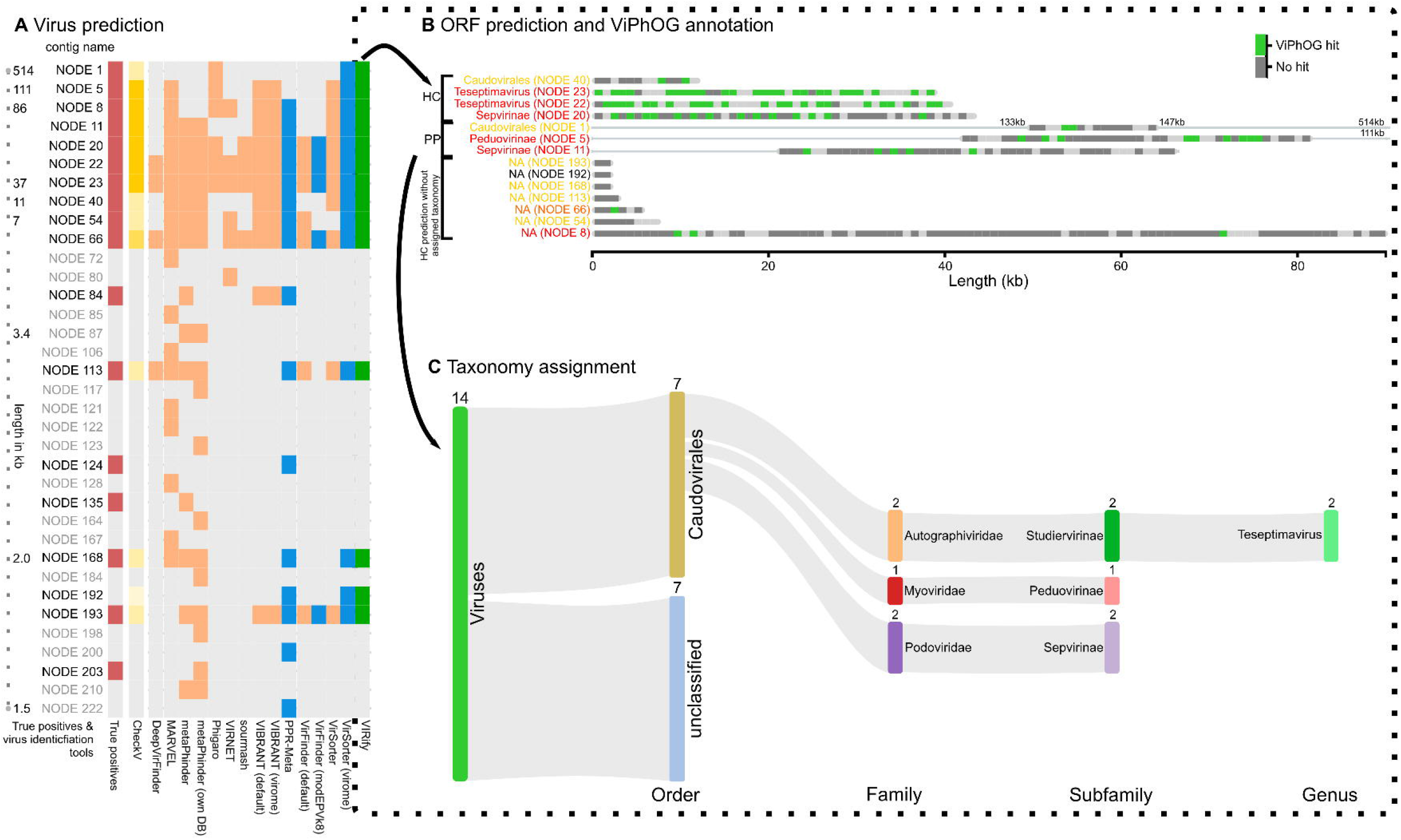
Viral contig selection and annotation procedure exemplar shown for the Kleiner co-assembly. **(A)** Comparison of virus predictions for the Kleiner co-assembly performed with various tools run via WtP. Shown are the 35 contigs (rows) predicted as viral by at least one of the tested tools. The column with red squares highlights the contigs manually identified as viral as described in the methods section. Blue squares indicate contigs predicted as viral by VirSorter (virome decontamination mode), VirFinder (VF.modEPV_k8.rda model) or PPR-Meta. The column with green squares indicates the viral contigs reported by the prediction approach implemented in VIRify, based on the results from WtP (see Methods section). Yellow squares indicate CheckV-quality results for contigs selected by VIRify that are either high-quality, medium-quality, low-quality or not-determined; going from dark (high-quality) to light yellow (not-determined). **(B)** ORFs predicted with Prodigal and annotated with the informative ViPhOGs for the 14 contigs identified as viral by VIRify. Of these, ten were predicted as high confidence (HC) and three as putative prophages (PP). No low confidence viral predictions were reported for the Kleiner co-assembly. The colored contig labels indicate the CheckV scores: red - high-quality, orange - medium-quality, yellow - low-quality, and black - not-determined. Grey bars indicate predicted ORFs without any ViPhOG hit while green bars indicate ViPhOG hits based on the model-specific bitscores. **(C)** Predicted viral sequences and corresponding taxonomic assignments based on informative ViPhOG hits for the Kleiner co-assembly.

For the Kleiner assembly, 17 contigs were identified as viral based on their alignment to the genome sequences of the phages that comprise the corresponding mock community (**Figure 3A**). VIRify identified 76.5% of the viral contigs in the Kleiner assembly and reported only one false positive prediction (NODE_192_length_1740_cov_2.233828), which also received no CheckV quality score. For the remaining contigs that VIRify predicted as viral, CheckV categorised six as high-quality, one as medium-quality, six as low-quality and the rest were indetermined (**Figure 3A**). All true negative predictions matched entries in NCBI’s RefSeq corresponding to *Salmonella* genomes, an observation consistent with the significant amount of *Salmonella* DNA that was previously detected in all of the viral-enriched samples used for obtaining the Kleiner assembly (Kleiner, Hooper, and Duerkop 2015). Regarding the reported false negatives, 3 corresponded to prophages predicted in the genome of *Salmonella enterica* subsp. *enterica* serovar Typhimurium str. LT2 and the remaining one matched entries in NCBI related to phage M13. According to the calculated F1-scores, the viral prediction tools that performed best for this assembly were VirSorter (using the virome decontamination mode) and VIRify, both of which had an F1-score of 0.84 (**SFigure 2**).

Regarding the Neto assembly, alignment of contigs to the genome sequences of the mock community viruses identified 259 of them as viral. In this case VIRify identified 51.7% of the viral contigs present in the assembly, and 21.6% of the contigs reported as viral were false positives. Among the false negatives, 75.2% of the contigs were derived from the genome of the mimivirus included in the mock community. In addition, 23.2% of the false negatives were contigs that ranged in size from 1,515 to 5,906 bp and that corresponded to prophages identified in the genomes of the *E. coli* and *B. thetaiotaomicron* strains included in the mock community. Based on the F1-scores, the tools that performed best for the Neto assembly were PPR-Meta (0.84), VirFinder with the VF.modEPV_k8.rda model (0.63) and VIRify (0.62) (**SFigure 2**). In total, VIRify predicted 170 contigs of the Neto co-assembly as viral, of which CheckV rated two as high-quality, three as medium-quality, 142 as low-quality, and 24 as indeterminate.

Overall, the calculated F1-scores revealed that VIRify and PPR-Meta were the tools that performed best for the prediction of viral contigs from the analysed mock community assemblies (**SFigure 2**). VIRify had a better performance than PPR-Meta on the Kleiner assembly, particularly due to its ability to predict prophages from assembled bacterial contigs (e.g. NODE_1_length_514496_cov_72.252612 and NODE_5_length_111535_cov_40.991774) and due to PPR-Meta being less precise than VIRify, as evidenced by the higher number of false positive predictions reported by the former (**Figure 3A**). On the other hand, PPR-Meta performed better than VIRify on the Neto assembly because the former tool had a higher recall rate that allowed the recovery of 87.6% of the mimivirus contigs, whereas VIRify recovered only 55.2% of the contigs derived from this virus.

#### Taxonomic classification of phage contigs and putative prophage regions down to the subfamily and genus ranks

The taxonomic annotation performed with VIRIfy on the Kleiner assembly classified seven of the putative viral contigs as members of the order *Caudovirales* (**Figure 3B-C**). Among them, two within the genus *Teseptimavirus* (genus), one *Peduovirinae* (subfamily), and two *Sepvirinae* (subfamily). Among the seven contigs with an assigned taxonomy, three were predicted by VirSorter to be putative prophage sequences (**Figure 3B**) which likely derived from bacterial contamination in the samples (Kleiner, Hooper, and Duerkop 2015). Of the six phages potentially included in the Kleiner co-assembly, VIRify identified contigs that correspond to phages P22 (NODE_20), T3 (NODE_23), and T7 (NODE_22). While contigs derived from phages T3 and T7 were correctly classified in the *Teseptimavirus* genus, the contig from phage P22 was incorrectly classified as a member of subfamily *Sepvirinae*. Nevertheless, both phage P22 and subfamily *Sepvirinae* belong to family *Podoviridae*, thus VIRify was able to classify the contig from phage P22 into the correct family. Overall, all the phages from the original mock community present among the putative viral contigs were correctly classified by VIRify, at least up to the family rank.

The pipeline reported a long 86 kb contig (NODE_8) from □VPE25 among the high confidence viral contigs, whose taxonomic lineage was not identified likely due to the low number of ViPhOG hits reported (**Figure 3B**). Phage M13 could not be identified either, as it was only assembled in small fragments due to low sequencing coverage, an observation that was consistent with the results reported by (Kleiner, Hooper, and Duerkop 2015). The M13 genome is 6.4 kb long and metaSPAdes only recovered two contigs in the high confidence set that matched different parts of the phage’s genome (NODE_113 with 2,682 bp and NODE_193 with 1,739 bp). Similarly, phage □6 was not even detected by (Kleiner, Hooper, and Duerkop 2015) after sequencing their viral-enriched samples, which explains why no contigs were identified for this phage in our coassembly.

#### Comprehensive detection of prokaryotic and eukaryotic viruses from a highly diverse mock-virome

Taxonomic annotation of the reported 170 putative viral contigs in the Neto assembly revealed that 65 of them were classified in the orders *Imitervirales* (58 contigs), *Caudovirales* (4), *Nidovirales* (2), and *Ortervirales* (1) (**Figure 4**). Among them, we found members of the virus families *Ackermannviridae, Myoviridae, Podoviridae, Mimiviridae, Coronaviridae*, and *Retroviridae*. For all these families, VIRify was also able to classify contigs at the subfamily level and some were also classified at the genus level: 49 contigs to *Mimivirus*, 2 contigs to *Alphacoronavirus*, and 1 contig to *Limestonevirus* (**Figure 4**). The taxonomic classifications provided by VIRify were compared with the taxonomy of the genomes from which the analysed contigs were derived. We determined as a result that VIRify’s taxonomic classification had an accuracy of 95.5% for the putative viral contigs detected in this assembly.

**Figure 4:**
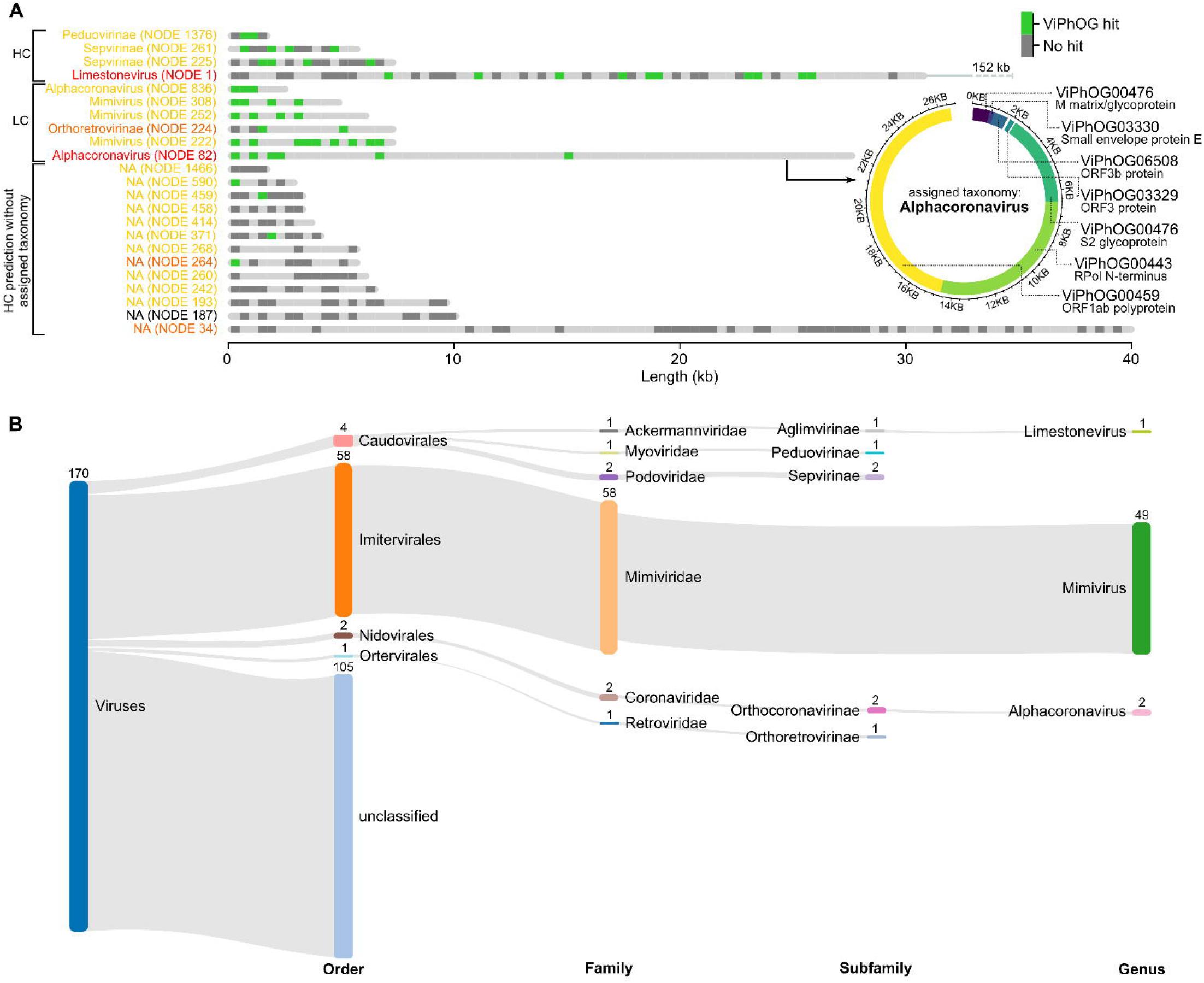
Predicted ORFs and corresponding ViPhOG annotations and taxonomy assignments for the Neto co-assembly. **(A)** ViPhOG-annotated ORFs for the Neto assembly. The colored contig labels indicate the CheckV scores: red - high-quality, orange - medium-quality, yellow - low-quality, and black - not-determined. VIRify assigned the genus *Alphacoronavirus* to NODE 82 based on seven informative ViPhOG model hits that are additionally shown as a circular visualization. NA - no taxonomy could be assigned due to missing model support. **(B)** Corresponding taxonomic assignments of Neto contigs based on the ViPhOG model hits shown in (A). Both visualizations can be automatically produced by the Nextflow version of VIRify and were only slightly manually adjusted via Inkcape.

Contigs classified as members of families *Myoviridae* and *Podoviridae* matched entries in NCBI’s nr database corresponding to phages of *E.coli* and *Bacteroides*, which suggests that these were most likely derived from DNA contamination by the bacterial component of the mock community. No contigs from the viruses that represented the families *Circoviridae, Polyomaviridae* and *Alphaflexiviridae* were detected among the length-filtered contigs by blastn, and therefore no contigs were assigned to any of the corresponding taxonomic lineages. Six contigs from viruses representing the families *Parvoviridae*, *Reoviridae* and *Herpesviridae* were detected among the set of low confidence viral contigs, although none of them were taxonomically classified by VIRify. The lengths of these contigs ranged from 1,543 to 2,356 bp and four of them corresponded to different segments of the segmented *Reoviridae* genome. Of note, one of these *Reoviridae* contigs contained a single putative protein that matched an informative ViPhOG of the genus *Rotavirus*, which corresponded to the taxonomic lineage of the *Reoviridae* virus present in the mock community. Furthermore, the contig from the *Herpesviridae* genome contained three putative proteins and one of them matched an informative ViPhOG of the subfamily *Alphaherpesvirinae*, which also agrees with the taxonomic lineage of the *Herpesviridae* virus present in the mock community. Nonetheless, the number of informative ViPhOG annotations in these contigs was insufficient to fulfil the criteria set for VIRify’s taxonomic assignment step.

### VIRify comprehensively reveals virus taxonomies in global ocean ecosystems

To showcase the utility of VIRify in uncovering viral diversity in ocean environments, we ran VIRify on 243 assemblies provided by the TARA Oceans project (Simon Roux et al. 2016; Gregory et al. 2019; Sunagawa et al. 2020), including 20 samples processed via 0.1-0.22 μm filters and thus potentially enriched for viral sequences (Gregory et al. 2019). As expected, we identified more complete/high-quality viruses in such samples enriched for smaller viruses in comparison to samples derived from larger 0.45 - 0.8 μm filters (**Figure 5A**). Across all assemblies and including high and low confidence viral predictions, VIRify identified viruses belonging to the order *Caudovirales* (n=395), *Imitervirales* (206), *Algavirales* (86), *Petitvirales* (9), and *Pimascovirales* (1). On the family level, the most prominent predictions were among *Autographiviridae* (360), *Myoviridae* (252), *Mimiviridae* (206), *Siphoviridae* (164), and *Phycodnaviridae* (86) (**SFigure 4**). In addition, we saw differences in the amount of predicted viral contigs between samples obtained from the same location but with DNA extracted using different filter sizes. For example, in CEUO01 (TARA_124_MIX) filtered for 0.1 - 0.22 μm (SAMEA2622799) 163 contigs were assigned to the order *Caudovirales*, whereas for the same sample material, but filtered for 0.45 - 0.8 μm (SAMEA2622801), only 15 *Caudovirales* contigs were found (**Figure 5A**). Interestingly, members of the genus *Prasinovirus* (large doublestranded DNA viruses that belong to the order *Algavirales*) were predominantly found in low confidence sets of 0.22-3 μm filtered fractions, which suggest that these viruses would have been missed if only VirSorter had been used on the data (**Figure 5B**).

**Figure 5:**
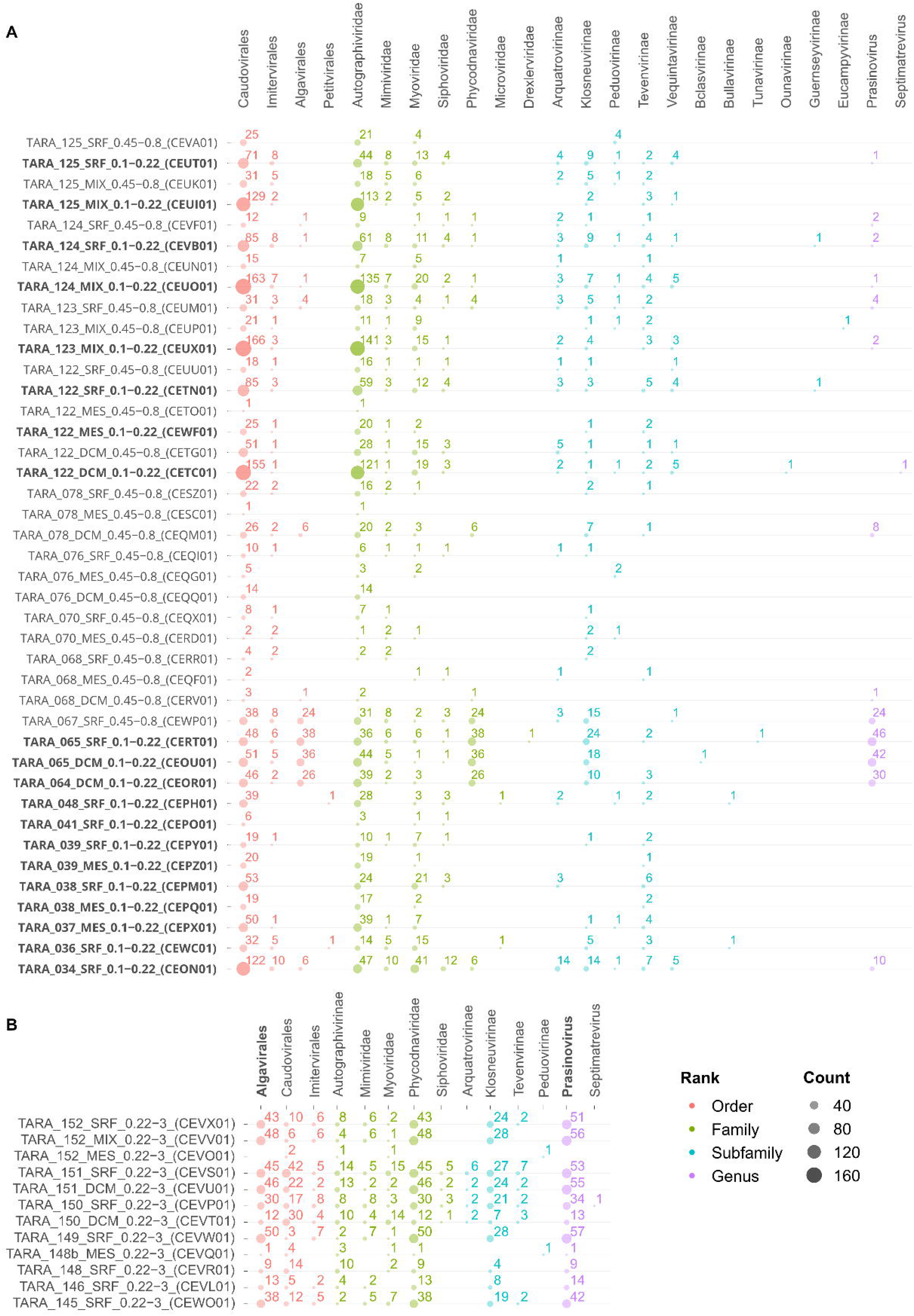
Viruses predicted and annotated by VIRify for 243 TARA Oceans assemblies. The assembly identifiers include information about the size fractionation of the corresponding sample. For example, samples obtained by smaller filter size 0.1-0.22 μm (and so expected to be enriched for smaller viruses) are labelled with the suffix _0.1-0.22. **(A)** Shows a selection (filters 0.1-0.22 μm and 0.45-0.8 μm) of 41 samples and the number of predicted viruses based on high confidence (VirSorter categories 1 and 2) and low confidence (VirSorter category 3 and combined VirFinder and PPR-Meta results) hits. More viruses are found for smaller filter sizes, as expected. Assemblies based on smaller filter sizes are highlighted in bold. For visualization purposes, we summarize high and low confidence predictions for samples labeled with the filter sizes 0.1-0.22 and 0.45-0.8. **(B)** Selection of 0.22-3 μm filtered samples with a high number of predicted prasinoviruses, large double-stranded DNA viruses belonging to the order *Algavirales*. These viruses are predominantly found in the low confidence set, thus they would be missed if only VirSorter were run on the data, but are predicted by our combination of VirFinder and PPR-Meta.

### ViPhOG models comprehensively cover viral proteins

Our comparison of virus-specific protein profile HMM databases included in VIRify showed that the ViPhOG models covered a large proportion of potentially viral CDS in agreement with the other databases (**Figure 6A**). In total, we predicted 1,204,455 CDSs from all high confidence viral contigs that VIRify reported for the 243 TARA assemblies, of which the ViPhOGs covered 56.7% (49.4% with applied bit score threshold) and thus were only slightly outperformed by the VOGdb (56.8%). The largest set of shared annotations comprised 284,791 CDS annotated as viral based on models from all compared databases except the RVDB (**Figure 6A**). In addition, the RVDB had the least predictions with only 15.3% annotated CDS in comparison to all other databases. Our model-specific bit score filtering reduces the amount of CDS annotations derived from ViPhOGs by 7.3% (87,869 CDS) (**Figure 6A**). While this might lead to the loss of potentially informative annotations that could be used for taxonomy assignment, we also remove false positive model hits, as shown in **Figure 6B**. Interestingly, the VPF derived from the IMG/VR database that also includes novel viral sequences derived from metagenome approaches, comprises a large proportion of unique models that match 117,962 CDS (9.8%) that are not covered by any of the other annotated databases. On the other hand, a significant number of CDS are annotated by models from the other databases but missed by the VPF models.

**Figure 6:**
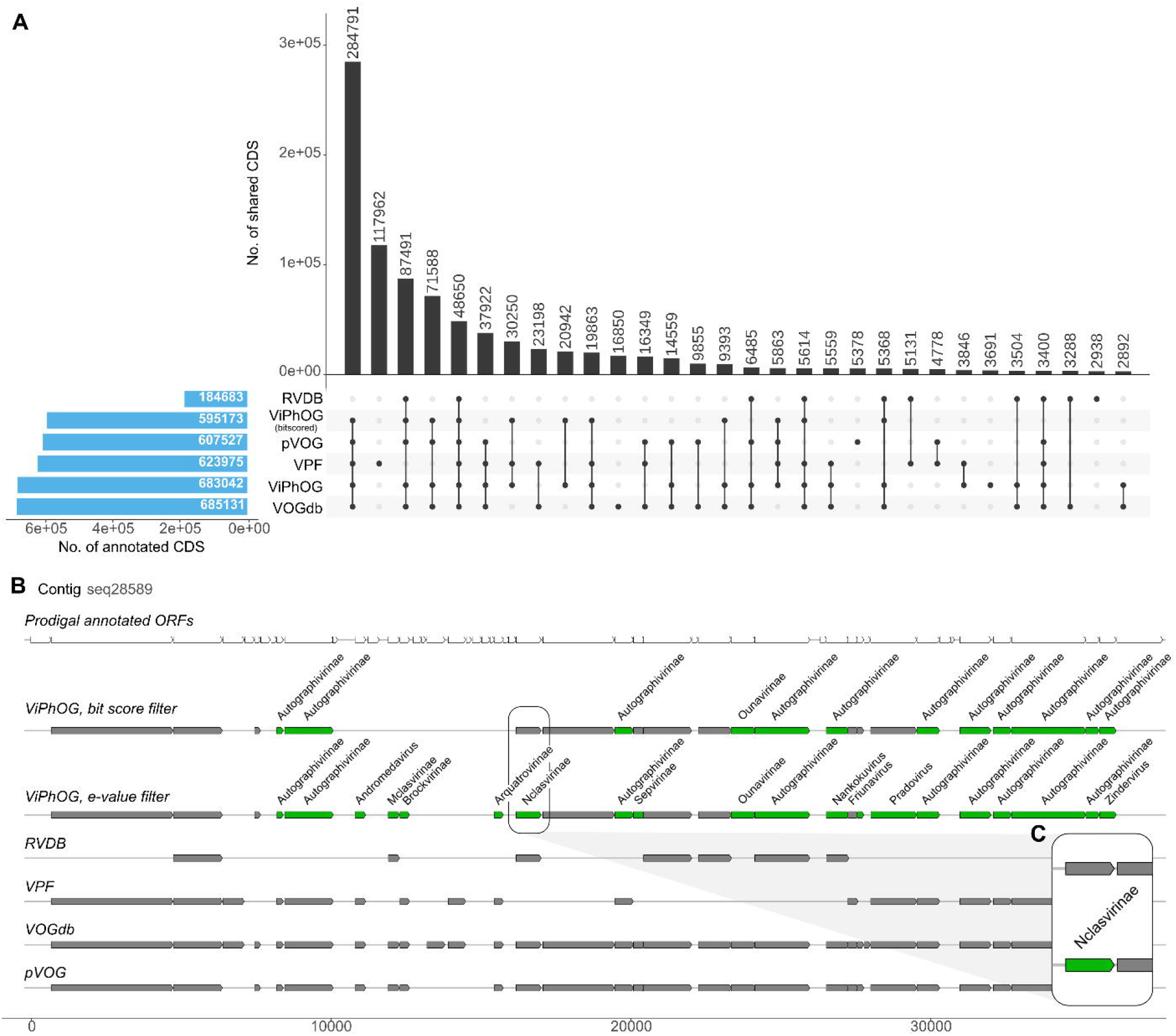
**(A)** Comparison of annotated CDS from VIRify’s high-confidence viral predictions from all 243 TARA Oceans assemblies. Our comparison shows that the ViPhOG models comprehensively cover a large proportion of potentially viral CDS in agreement with other public databases. The RVDB had the fewest predictions with only 15.3% annotated CDS. Results of the VPF database, where models are derived from the IMG/VR database that also includes novel viral sequences derived from metagenome approaches, comprises a large proportion of unique models that exclusively match 117,962 CDS (9.8%) that are not covered by any of the other databases. However, a significant number of CDS are annotated by models from the other databases but missed by VPF. Our model-specific bit score filtering (ViPhOG - threshold) reduces the number of CDS annotations derived from ViPhOGs by 7.3% **(B)** Visualization of predicted (top row) and annotated ORFs for one exemplar contig from TARA Oceans assembly CEUI01. Grey bars indicate hits against an HMM of the corresponding database, while informative ViPhOG hits with taxonomic information are shown in green. The top ViPhOG track shows hits filtered by bit score (or e-value if no bit score could be assigned for a model, see Methods) and the bottom ViPhOG track shows e-value-filtered hits. **(C)** While the model-specific ViPhOG bit score threshold can lead to the loss of potentially informative annotations for taxonomy assignment, it can also reduce the number of false positive model hits and thus increase the overall accuracy of VIRify.

## Discussion

Metagenomic surveys of different environments during the last few decades have had a profound impact on the rate at which novel viruses are being discovered (Call, Nayfach, and Kyrpides 2021). Despite the resulting sharp increase in the number of publicly available viral genomes, viruses remain relatively understudied in many environments and the taxonomic profiling of viral communities is still challenging due to the lack of universal genetic markers that support the phylogenetic resolution of viral taxa (Pratama and van Elsas 2018; Dion, Oechslin, and Moineau 2020). As a contribution to the currently available repertoire of tools designed to address these challenges, we designed and implemented the VIRify pipeline for detecting and taxonomically annotating viral contigs in metagenomic datasets.

The results presented here for the analysis of the mock community assemblies with VIRify demonstrated that it is a suitable pipeline for generating highly accurate taxonomic profiling of viral communities present in metagenomic datasets. In comparison with most of the tools currently used for detecting viral contigs in metagenomic assemblies, VIRify demonstrated higher predictive performance for contigs ≥ 1.5 kb in both the Keiner and Neto assemblies (**Figure 3A**, **SFigure 2** and **SFigure 3**). Furthermore, the use of our manually-curated informative ViPhOGs led to the taxonomic classification of putative viral contigs from the Neto assembly with an accuracy of 95.5%. Regarding the Kleiner assembly, all classified putative viral contigs were assigned to the correct taxonomic lineages at different degrees of resolution, with some classified up to the family rank and other classified up to the genus rank. The pipeline also provided taxonomic classifications for putative prophages identified in contaminant bacterial contigs from both mock assemblies.

VIRify’s viral prediction performance was the result of combining the effectiveness of VirSorter in predicting phage and prophage sequences (Simon Roux, Enault, et al. 2015; Simon Roux, Hallam, et al. 2015), with the ability of PPR-Meta and VirFinder’s VF.modEPV_k8.rda model to predict sequences from eukaryotic viruses, as evidenced by their F1-scores for the Neto assembly (**SFigure 2**). Overall, most of the viral prediction tools evaluated here demonstrated a low performance for the Neto assembly, which indicates that these tools were mainly developed for the detection of phages and are not suited for the comprehensive detection of eukaryotic viral sequences. Considering that PPR-Meta was originally designed for the detection of phages and plasmids (Fang et al. 2019) and that the Neto assembly mainly comprised eukaryotic viruses, it was striking to find that PPR-Meta was the tool that performed best for this assembly. Despite this observation, we decided to keep VIRify’s original viral prediction approach due to the fact that VirSorter predictions include the identification of prophages, and that in terms of precision VIRify performed as well or better than PPR-Meta for the analysed mock community assemblies. In addition, a recent study by (Lai et al. 2020) used the same three tools selected for VIRify to successfully identify ~400,000 phages from 66,425 human metagenomic samples from a diverse range of body sites.

Several studies have reported collections of profile HMMs for groups of orthologous viral genes, such as pVOGs (Grazziotin, Koonin, and Kristensen 2017), vFAMs (Skewes-Cox et al. 2014), RVDB (Bigot et al. 2019) and VOGdb (http://vogdb.org/). The pVOGs database have been previously applied for the taxonomic classification of phages in human gut metagenomes (Waller et al. 2014), and VOGdb includes data on the lowest common ancestor associated with each profile HMM. However, unlike VOGdb, the informative ViPhOGs we curated have bespoke bit score thresholds that were carefully selected to decrease the likelihood of false positive taxonomic assignments. As illustrated in **Figure 6B**, using the informative ViPhOGs with a single e-value threshold resulted in a higher chance of false positive taxonomic annotations, as was the case for contig seq28589 from the assembly of the TARA Oceans sample CEUI01 (study accession PRJEB7988). Therefore, our results indicate that the use of curated bit score thresholds offers an adequate balance between predictive power and classification accuracy. This strategy has been successfully implemented previously in Pfam and TIGRFAMs, some of the most widely used protein family databases for the functional annotation of genomes and metagenomes (Mistry et al. 2021; Haft et al. 2001).

We further showed that VIRify can be applied to metagenome-assembled genomes obtained from large environmental studies by using 243 TARA Oceans assemblies (Simon Roux et al. 2016; Gregory et al. 2019; Sunagawa et al. 2020), including samples enriched for viral sequences (Gregory et al. 2019). As expected, we identified more viruses in enriched samples that used smaller filter sizes in comparison to samples derived from larger filters (**Figure 5A**). In accordance with (Gregory et al. 2019), a high proportion of predicted viruses could not be taxonomically classified to a known viral family. Successfully classified contigs were predominantly assigned to dsDNA viral families and phages. Interestingly, prasinoviruses were predominantly found in low confidence sets, thus, would have been missed by only running VirSorter on the data (**Figure 5B**). Here, our combination of VirFinder and PPR-Meta predictions helped to recover contigs that the pipeline could later taxonomically assign to this genus. Due to their larger size, prasinoviruses are underrepresented in smaller size fractionated samples, underlying the importance of appropriate filtering and enrichment steps and their combination to comprehensively collect viruses from environmental samples.

Despite the great performance observed for the analysed assemblies, the results obtained here revealed a few limitations of the current VIRify pipeline. As evidenced by analysis of the Neto assembly, VIRify’s ability to detect contigs from eukaryotic viruses is currently suboptimal. Increasing the pipeline’s sensitivity will require the careful evaluation of novel viral prediction tools that could easily be incorporated or used to replace the ones currently used. For example, VirSorter might be replaced by the recently released VirSorter2 (Guo et al. 2021) in a future version of VIRify. An additional limitation of VIRify is the existing bias in the extent to which the informative ViPhOGs represent the different genera in the current viral taxonomy. Phages that belong to the *Caudovirales* represent 94% of the phage sequences currently available in NCBI’s viral RefSeq, which indicates that the great majority of phage isolates characterized to date are members of the *Caudovirales*. Nevertheless, the analysis of viromes from different environments has led to a significant increase in the number of sequences from uncultivated viruses, which have contributed to increasing the available information on underrepresented viral taxa and the identification of novel ones. Including data on these uncultivated viruses, such as those available in the IMG/VR database (David Paez-Espino et al. 2019), will certainly help balance the coverage of the whole viral diversity by the set of informative ViPhOGs used within the VIRify pipeline. Finally, the quality of the metagenomic assembly is key for VIRify’s performance due to the challenge that short viral contigs pose on their detection by the viral prediction tools currently used in the pipeline (**Figure 3A**). Furthermore, short contigs will generally contain fewer complete CDS, and thus will be less likely to have the minimum number of ViPhOG hits required by the pipeline to provide a taxonomic assignment.

Although VIRify has been benchmarked and validated with metagenomic data in mind, it is also possible to use the pipeline to detect RNA viruses in metatranscriptome assemblies (e.g. SARS-CoV-2, see Hufsky et al. 2021). Some additional considerations in this regard include 1) quality control, 2) assembly, 3) post-processing, and 4) classification. Some brief recommendations on how to prepare metatranscriptome assemblies and run them through VIRify can be found in the GitHub manual.

Overall, our results demonstrated the utility of VIRify for the analysis of viral communities in metagenomic datasets. In contrast with other resources currently available, VIRify offers an enhanced capability to predict contigs from eukaryotic viruses and the ability to taxonomically classify viral contigs using a set of carefully curated HMMs and their bespoke bitscore thresholds. Future versions of VIRify will attempt to improve even further the sensitivity of viral contig prediction and to extend the ViPhOGs’ taxonomic coverage by including sequences from uncultured viruses. In addition, the taxonomic classification of contigs from small viruses could be improved by adjusting the taxonomic voting system to accommodate differences in the number of available informative ViPhOGs and in the average genome size observed between different viral lineages. VIRify will be fully integrated into the MGnify suite of metagenomic analyses hosted at the EMBL-EBI, which is continuously supported and updated to suit the needs of the scientific community.

## Acknowledgements

MH was supported within the Collaborative Research Centre AquaDiva (CRC 1076 AquaDiva) of the Friedrich Schiller University Jena funded by the Deutsche Forschungsgemeinschaft and additionally appreciates the support of the Joachim Herz Foundation by the add on fellowship for interdisciplinary life science. GRP was supported by BB/P027849/1 (CABANA project for capacity building for bioinformatics in Latin America). The authors thank Fransiska Hufsky for improving the readability of the illustrations.

## Supplementary Figure captions

**SFigure 1:** Informative ViPhOGs’ coverage of viral taxonomy. Circular dendrogram showing the NCBI virus taxonomy for all currently known viral genera, with taxa covered by informative ViPhOGs highlighted with either dots or bars, and using the same colour scheme as in **Fig. 2**. For the taxonomic ranks (order, family and subfamily) dots were used to indicate whether any informative ViPhOGs were identified for the corresponding taxon. By contrast, bars were used for taxa in the genus rank (tree’s leaves) to indicate the number of different informative ViPhOGs identified for the corresponding genus, as indicated by the numeric scale in the outer rings.

**SFigure 2:** F-score comparison for virus prediction results from the Kleiner and Neto assemblies for all tools run via WtP and VIRify (combination of VirSorter, VirFinder with VF.modEPV_k8.rda model and PPR-Meta).

**SFigure 3:** UpSet output for the Neto assembly calculated with WtP. Viral prediction tools are compared and overlapping sets are shown.

**SFigure 4:** Abundance plot of viral ranks predicted for 243 TARA Oceans assemblies (https://www.ebi.ac.uk/ena/browser/view/PRJEB22493) using the VIRify pipeline. Combined results for high confidence, low confidence, and putative prophage hits are shown. VIRify was run in v0.2 and with the --virome option and a contig length filtering of 5000 nt.

## Notes

https://doi.org/10.17605/OSF.IO/FBRXY

### Competing Interest Statement

The authors have declared no competing interest.

