## Supplementary figures and images for "VIRify: an integrated detection, annotation and taxonomic classification pipeline using virus-specific protein profile hidden Markov models"

### Supplemental Figure 1

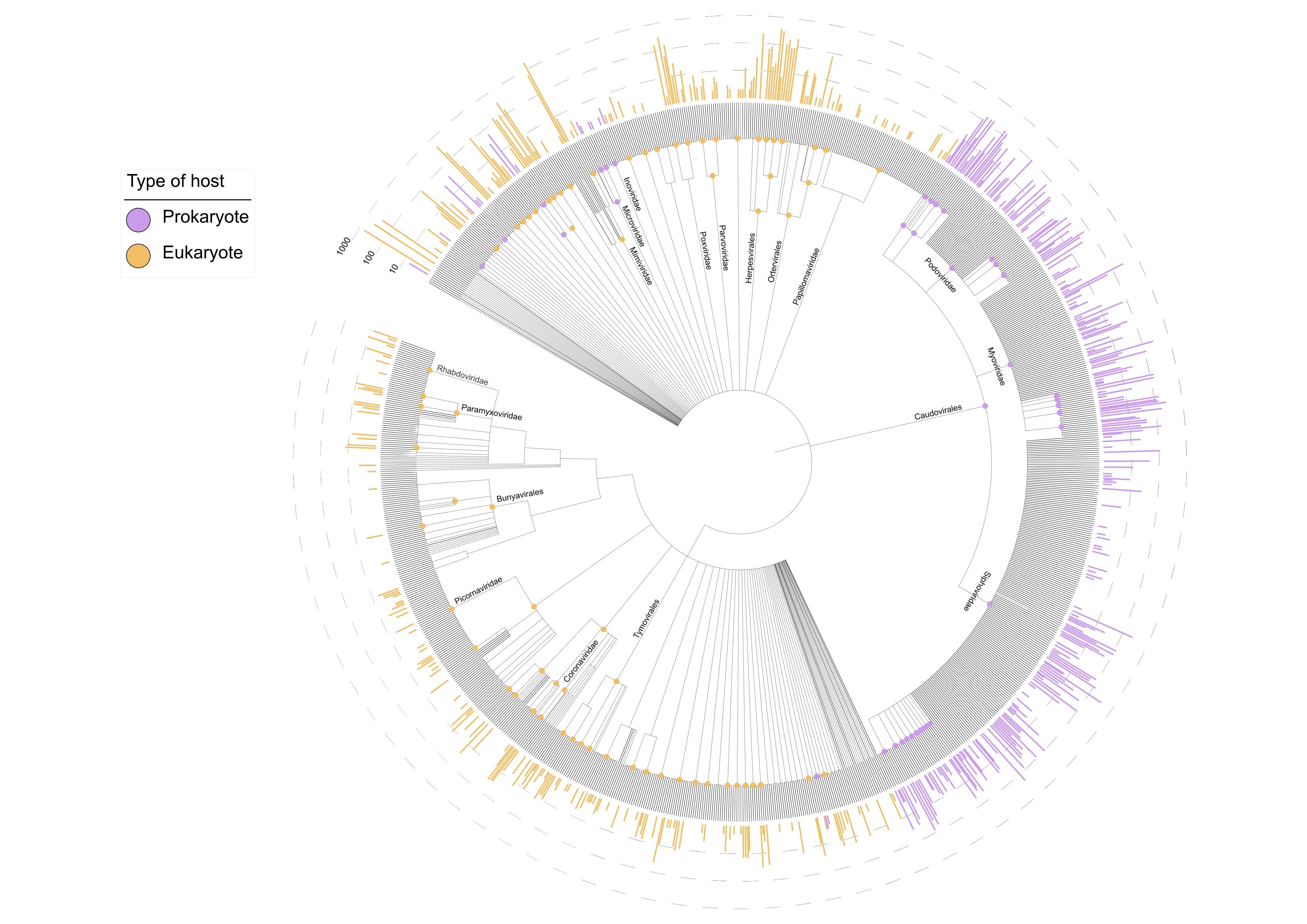

### Supplemental Figure 2

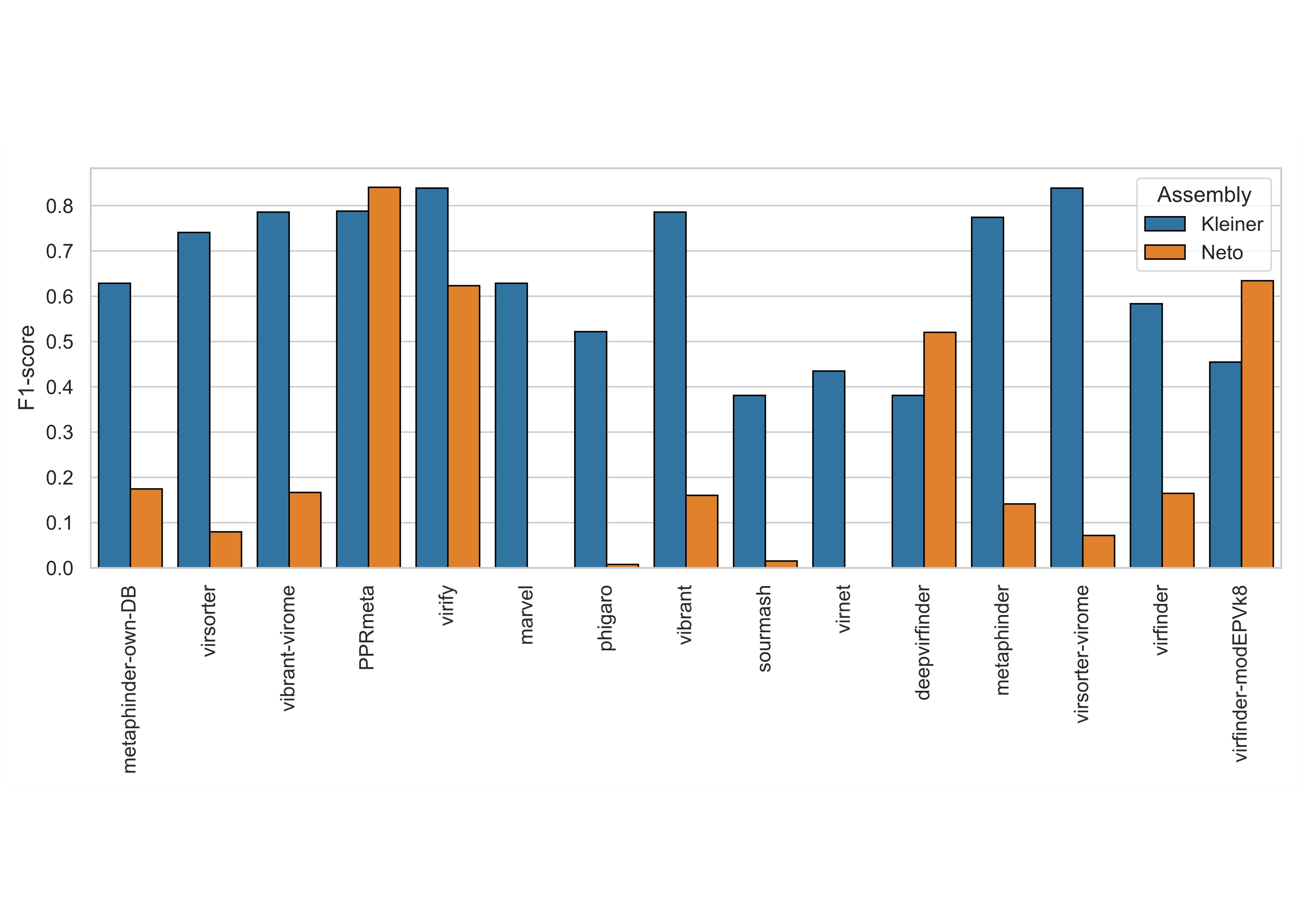

### Supplemental Figure 3

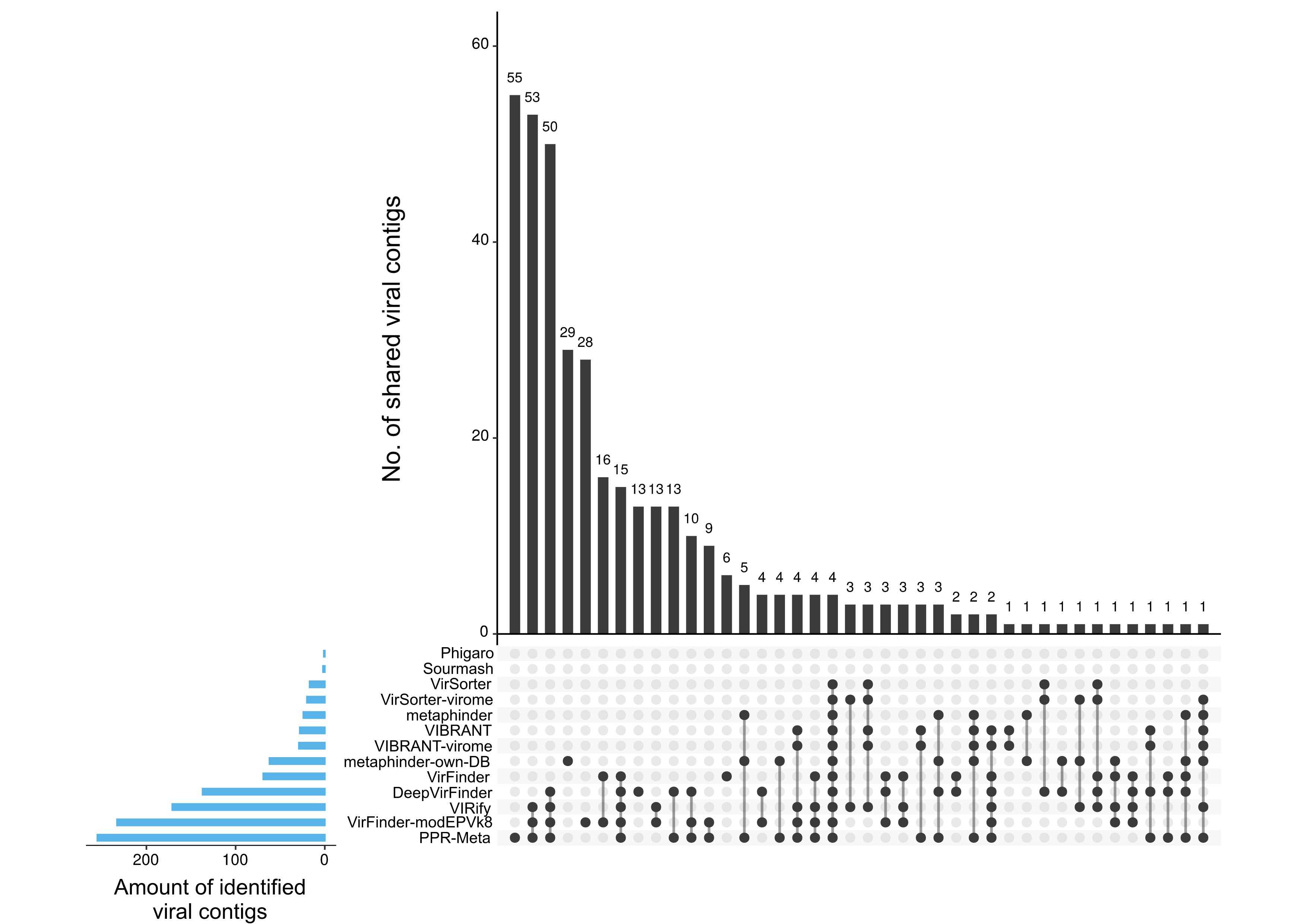

### Supplemental Figure 4

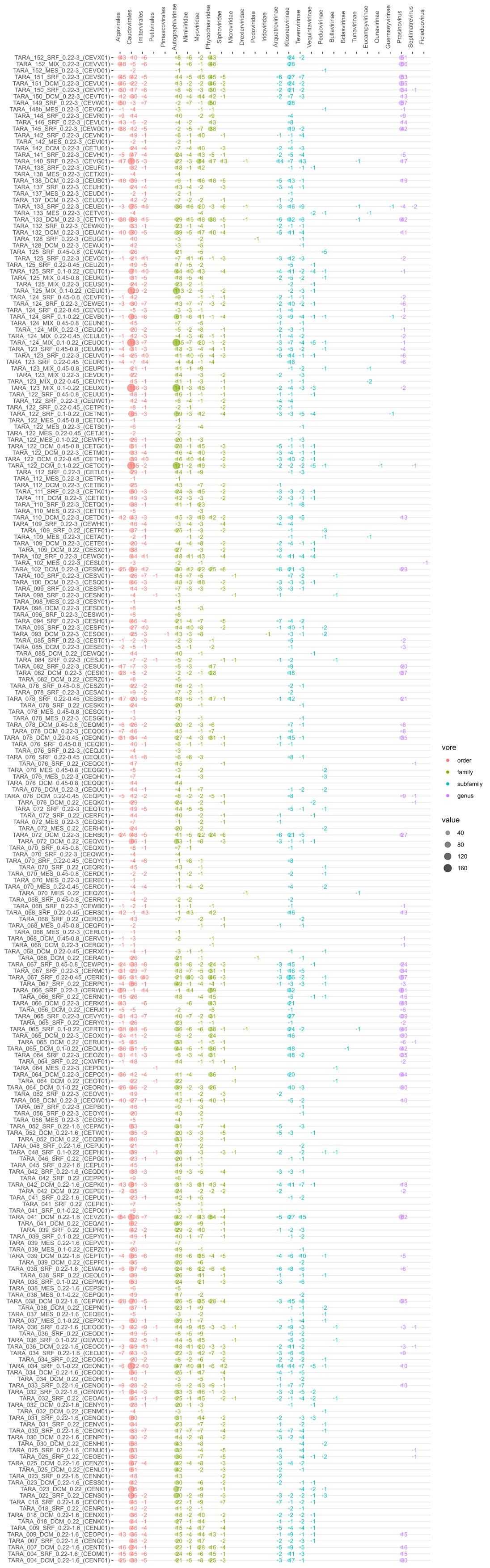
